# Neural correlates of modal displacement and discourse-updating under (un)certainty

**DOI:** 10.1101/2020.07.03.187112

**Authors:** Maxime Tulling, Ryan Law, Ailís Cournane, Liina Pylkkänen

**Affiliations:** Department of Linguistics, New York University, New York, NY 10003; NYU Abu Dhabi Institute, New York University Abu Dhabi, Abu Dhabi, United Arab Emirates; Department of Psychology, New York University, New York, NY 10003

## Abstract

A hallmark of human thought is the ability to think about not just the actual world, but also about alternative ways the world could be. One way to study this contrast is through language. Language has grammatical devices for expressing possibilities and necessities, such as the words *might* or *must*. With these devices, called “modal expressions,” we can study the actual vs. possible contrast in a highly controlled way. While factual utterances such as “There is a monster under my bed” update the *here-and-now* of a discourse model, a modal version of this sentence, “There might be a monster under my bed,” displaces from the *here-and-now* and merely postulates a possibility. We used magnetoencephalography (MEG) to test whether the processes of discourse updating and modal displacement dissociate in the brain. Factual and modal utterances were embedded in short narratives, and across two experiments, factual expressions increased the measured activity over modal expressions. However, the localization of the increase appeared to depend on perspective: signal localizing in right temporo-parietal areas increased when updating others’ beliefs, while frontal medial areas seem sensitive to updating one’s own beliefs. The presence of modal displacement did not elevate MEG signal strength in any of our analyses. In sum, this study identifies potential neural signatures of the process by which facts get added to our mental representation of the world.

## INTRODUCTION

Speculating about possibilities employs our unique human capacity to displace from the *here-and-now* (Hockett, 1959; Bickerton, 2008; Suddendorf et al., 2009). We can express possibility using ‘modal expressions’ like “There *might* be a monster”, shifting our perspective from the immediate present to a hypothetical scenario. Other cognitive abilities that shift into alternative perspectives, like thinking about the past or future and conceiving the viewpoints of others, seem to share a brain network consisting of hippocampal and parietal lobe regions (Buckner & Carroll, 2007; Mullally & Maguire, 2014). However, we know surprisingly little about the neural mechanisms involved in modal displacement. While factual statements like “There is a monster” update our beliefs about a situation, modal utterances indicate uncertainty instead. Are the mental operations of discourse updating and modal displacement dissociable in the brain? Here, we investigated the neural correlates of integrating factual and modal utterances into an existing discourse representation.

### Cognitive Processes Involved with Comprehending Discourse

When comprehending discourse, we represent the perspective, place and time of the discussed situation (van Dijk & Kintsch, 1983; Zwaan & Radvansky, 1998), and distinguish between facts and possibilities compatible with the *here-and-now* of this alternative reality. Consider this scene from Ovid’s tale about the ill-fated lovers Pyramus and Thisbe.

> *When a lioness, bloody from hunting, approaches, Thisbe flees into a cave, losing her shawl in the process. As Pyramus encounters the lioness hovering over Thisbe’s bloodstained shawl with his lover nowhere in sight, he quickly concludes she **must** have been devoured by the beast.*

All but the underlined sentence are factual claims made about the actual state of affairs (Stalnaker, 1996). We use these utterances to build a mental situation model, which is dynamically updated as new information becomes available (Glenberg et al., 1987; Morrow et al., 1989; Zwaan & Madden, 2004). Maintaining these discourse models elicits activation in the medial prefrontal cortex (mPFC), posterior cingulate cortex (PCC) and temporo-parietal areas (Speer et al., 2007; Whitney et al., 2009; Xu et al., 2005; Yarkoni et al., 2008). To interpret the narrative above, we also engage in higher order cognitive processes such as modal displacement and Theory of Mind (ToM) reasoning (Premack & Woodruff, 1978). ToM is the ability to represent someone else’s belief state separately from our own, allowing us to understand how Pyramus induced that Thisbe died, even though we know she is still alive. Pyramus based his conclusion on indirect evidence (the bloody shawl), signaling with the modal verb *must* that the devouring is not actual or known. Modals like *must* or *may* allow reasoning about open possibilities compatible with a situation (Kratzer, 1981, 2012; Phillips & Knobe, 2018; von Fintel, 2006).

Since ToM and modal displacement both require a representation that is different from the actual situation (Phillips & Norby, 2019), they may recruit overlapping brain areas. While, there has been no systematic study of the neural bases of modal processing, ToM tasks are consistently reported to activate the dorsal/posterior inferior parietal lobule (IPL), temporoparietal junction (TPJ), medial prefrontal cortex (mPFC), posterior cingulate cortex (PCC) and rostral anterior cingulate cortex (rACC) (e.g. Koster-Hale et al., 2017; Mahy et al., 2014; Schurz & Perner, 2015). In particular the right TPJ seems involved in representing other’s mental state (Saxe & Powell, 2006; Saxe & Wexler, 2005; Vistoli et al., 2011) though some suggest this activity may be attributable to more domain general cognitive processes such as reorienting attention (Corbetta et al., 2008; Decety & Lamm, 2007; Mitchell, 2008; Rothmayr et al., 2011).

### This Study

How do our brains distinguish between information that states facts versus information that only conveys possibilities? We investigated the differences between factual and modal language comprehension in two experiments (Figure 1). We used magnetoencephalography (MEG), providing us with high temporal resolution and relatively good spatial localization of brain activity during sentence comprehension. Experiment 1 investigated the neural bases of discourse updating and modal displacement by contrasting sentences that contain modal verbs against sentences containing the factual verb ‘do’ embedded in short narratives. In experiment 2, we further investigated under which conditions discourse updating takes place by manipulating the certainty of the sentential context in which the target verbs (factual vs. modal) were embedded: factual (certain), conditional (uncertain) or presupposed (already known). Discourse updating should take place under actual situational changes (e.g. when new factual information is added to a factual context), but not when novel information is hypothetical (modal conditions) or when the entire context is hypothetical (conditional context). Modal displacement should occur whenever utterances postulate hypothetical possibilities.

**Figure 1.**
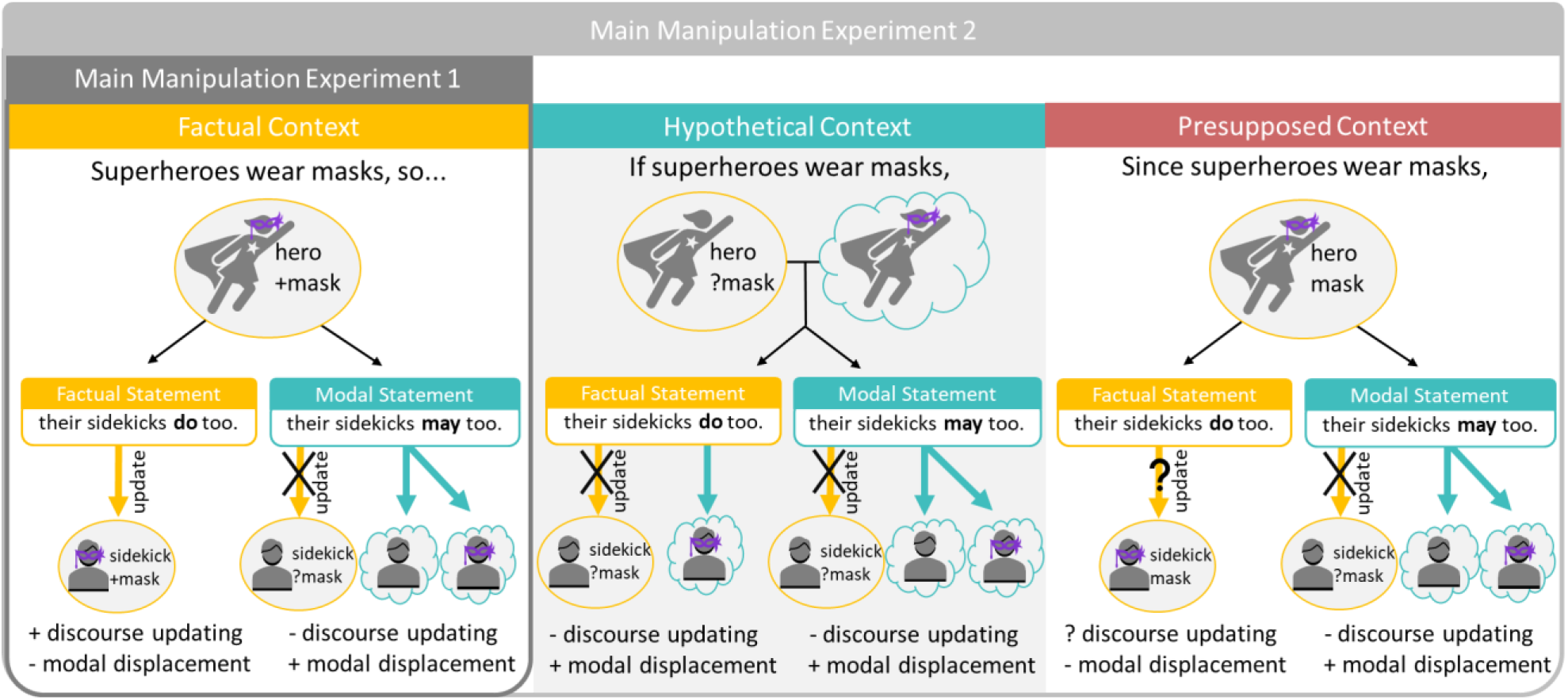
Simplified Illustration of Main Manipulations Experiment 1 and 2. Model of operations assumed to be present during the processing of factual (yellow) and modal (teal) statements (simplified from actual stimuli). Experiment 1 contrasts factual and modal statements in a factual discourse context, while Experiment 2 varies whether the discourse context is factual, hypothetical, or presupposed. Updating of the discourse situation model (round) is expected to take place under certainty (in factual contexts with a factual update). Both modal (*may*) and conditional expressions (*if superheroes wear masks*) evoke hypothetical situations (cloud) involving modal displacement. Since the presupposed context marks information already known, we are not sure whether updating would take place.

## METHODS

### Experiment 1

#### Participants

26 right-handed, native English speakers participated in the experiment (4 male) taking place at the New York University (NY) campus. One participant was excluded from further analysis for having an accuracy lower than 70% on the behavioral task. The age range of the remaining 25 participants was 19-52 years old (M= 25.7, SD = 7.46). All participants had normal or corrected to normal vision, no history of neurological impairment and provided informed written consent.

#### Stimuli

We developed an experimental paradigm where we contrasted the modal verbs *may* and *must* against the factual auxiliary verb *do*. In order to have *do* naturally appear in the same position as *may* and *must*, our sentences contained verb phrase (VP) ellipsis, e.g. “Normally only knights <u>sit at the round table</u>, but the king says that the squires *may*/*must*/*do* <sit at the round table> too.” While the verb *do* indicates factuality, modals indicate hypothetical scenarios that are compatible with the actual world given someone’s knowledge or the set of circumstances. We specifically chose to use the modal expressions *may* and *must* because they vary among two dimensions: ‘modal force’ and ‘modal base’. Modal force refers to the likelihood of a hypothetical situation, i.e. whether it is deemed a possibility (*may*) or a necessity (*must*). The modal base denotes what we base this likelihood assessment on: our knowledge or the circumstances, e.g. rules/norms. The modals *may* and *must* are ambiguous in allowing for both a knowledge-based (e.g. “Given what I know, there may/must be a monster under my bed”) and a rule-based reading (e.g. “Given what the rules are, you may/must eat your dinner now”). Using such ambiguous modals, we could compare the effect of modal base without varying the form of the target item.

We constructed 40 sets of short English narratives. Each story consisted of three sentences, starting with a context sentence designed to either bias towards a knowledge-based (epistemic) scenario, or a rule-based (deontic) scenario. The context sentence was followed by a target sentence and each story ended with a final task sentence that was either congruent or incongruent with the previous two sentences (Figure 2A). The target sentences contained the target modal verb (the possibility verb *may* or the necessity verb *must*) and were compared against the factual condition containing the verb *do*. In the context sentence a property or habit was introduced that applied to one group (e.g. “knights sit at the round table”), and the target sentence indicated this was also (possibly) the case for another group (e.g. *their squires do/may/must too*). Each stimulus set therefore consisted of 6 sentences (2×3, BASE: [knowledge, rules] × FORCE: [possibility, necessity, factual]) adding up to a total of 240 sentences for all 40 stimuli sets (Figure 2B). The third sentence of the story was a task sentence either congruent (50%) or incongruent (50%) with the prior two sentences. One third of the task sentences were specifically tapping into the congruency of the modal base (Figure 2C). Across conditions, how often task items were congruent or incongruent with the preceding sentences was controlled for, as was how often questions tapped into information obtained from the context or target sentence.

**Figure 2.**
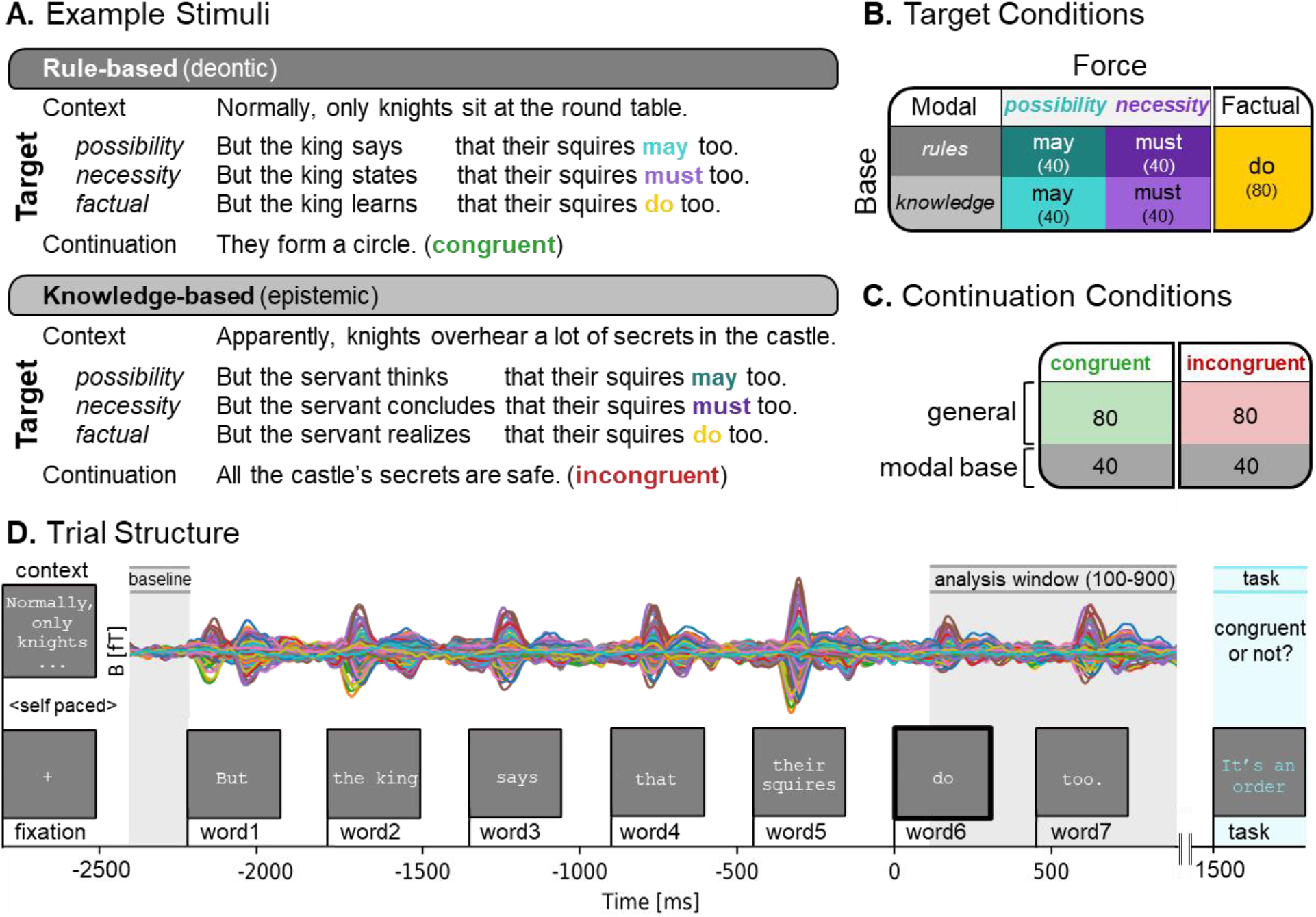
Design and procedure Experiment 1. **A**: Example stimuli set. Short narratives consisted of three parts. A context sentence biasing towards a rule-based or knowledge-based modal interpretation, followed by the target sentence containing one of the target verbs varying in force (possibility, necessity or factual). The third continuation sentence was either congruent or incongruent with prior sentences. Details on controlled between-stimuli variation can be found in Appendix I. **B**: Experimental design with number of items per condition in brackets (total = 240). The stimuli vary along two dimensions: MODAL BASE [rules, knowledge] and FORCE [possibility, necessity, factual]. **C**: Continuation Conditions. Half of the continuations are incongruent with the previous sentences. One third tap into modality and are congruent or incongruent with the modal base of the previous sentences. **D**: Trial structure with evoked MEG responses from one participant. A context sentence was displayed until participants pressed a button. After a fixation cross (300ms) the target sentence was displayed word-by-word for 300ms each followed by a 150 ms blank screen. The continuation sentence was displayed with a 600ms delay, and participants indicated by button press whether this was congruent or incongruent with the prior story. Time windows for baseline correction (−2450 to −2250ms) and statistiacal analysis (100-900ms) are relative to the target verb (word6) onset.

All target sentences had the same sentence structure: CONNECTIVE (but/and/so)| the | NOUN.SG | VERB1 | that | DETERMINER | NOUN.PL | TARGET (may/must/do) | <ELIDED VP> too. The embedded clause of the sentence (introduced by *that*) was kept consistent across all conditions. We controlled for between-item variation in the other parts of the stimuli along the following dimensions: the count of different CONNECTIVES and DETERMINERS among the modal base conditions, the average length, frequency, number of syllables and morphemes of NOUN.SG among different modal base conditions, and the average length (in words and letters), stativity, transitivity and structural complexity of the <ELIDED VP> material in the target sentence across different base conditions (see Appendix I). The information on lexical frequency and morpheme length was obtained from the English Lexicon Project (Balota et al., 2007). Within the modal base dimension, the target sentences only varied in the embedding verb (VERB1) to support biasing the reading of the target modal verb. Embedding verbs were divided into three categories occurring with knowledge-based, rule-based or factual targets. Each verb category contained 12 different verbs, which were repeated maximally 7 times across the entire experiment. Between the two base conditions, the knowledge-based and rule-based sentences also differed in their preceding context sentence and subject, to help bias the interpretation of the ambiguous modals *may* and *must*. In order to encourage the rule-based reading, the context introduced an event that was compatible with both permission or obligation (e.g. sitting at the royal table), and the target sentence introduced a subject that was in an authority position over the sentence object (e.g. a king over squires). In order to encourage the knowledge-based reading, the context introduced an event that was very unlikely to be permitted or obliged (e.g. overhearing secrets) and the target sentence introduced a subject that was in a bystander position to the event (e.g. a servant).

The effectiveness of the biasing conditions was tested with a survey on Amazon Mechanical Turk made with the help of Turktools (Erlewine & Kotek, 2016). For this norming, the target sentences containing modal verbs (160 items in total) were adjusted so that unambiguous adjectives replaced the ambiguous target modal verbs. Knowledge-based *may* was replaced with *are likely to*, knowledge-based *must* with *are certain to*, rule-based *may* with *are allowed* to and rule-based *must* with *are obliged* to. E.g. the target sentence “But the king says that the squires may too” became “But the king says that the squires are allowed to as well”. These unambiguous target sentences were then displayed with their preceding context sentence and a gap substituting the adjective. Participants (n=320) were asked to choose which of 4 options (*obliged, allowed, likely* and *certain*) would fit the gap best. Each target sentence was judged 32 times across all participants. The experiment took about 2-4 minutes and participants were paid $0.20 for completing the experiment. Each participant completed 25 sentences, comprised of 20 test items and 5 filler items that served as an attention control, in random order counterbalancing for condition. Results were excluded from participants that indicated to not have English as a native language (n=17) and from participants that made more than 1 mistake on the filler items (n=6). For the responses of the remaining 297 participants we noted whether the modal base of their response (*allowed* and *obliged =* rule-based, *likely* and *certain* = knowledge-based) matched the intended modal base of the target items or not. For each item, we calculated the average percentage of matches with the intended modal base (bias score), and only approved an item for the experiment if its bias score was 70% or higher. This norming happened in two parts. In the first round, all 160 items were tested, and 137 items were accepted. The remaining 23 items had a bias score below the 70% threshold and were altered to improve their bias. In the second round, these 23 items were re-tested (now mixed with a random selection of the previously approved items) and judged with the same criteria. This time 18 items were accepted, and 5 scored below the 70% threshold. The 5 items that did not pass the norming experiment were altered again with the help and approval of several native speakers, and then included into the experiment.

The lexical frequency of knowledge-based (epistemic) and rule-based (deontic) readings of *may* and *must* are not evenly distributed in written American English: the verb *may* is knowledge-based about 83% of the time (Collins, 2007), while *must* is knowledge-based 16% of the time (Hacquard & Wellwood, 2012), in all other cases the verb has a circumstantial base that includes rule-based meanings. While these lexical frequency differences may have an effect on the processing of the individual items, we expect that grouping the different levels of the force (grouping *knowledge-based* and *rule-based* responses together) or modal base manipulation (grouping *possibility* and *necessity* responses together) should wash out any effects of this imbalance.

#### Procedure

Before recording, the head shape of each participant was digitized using a FastSCAN laser scanner (Polhemus, VT, USA). Additionally, we recorded the location of three fiducial locations (the nasion, and left and right preauricular points) and five reference points for purposes of co-registration. Before participants entered the MEG-room they received verbal instructions and did a short practice block (of eight trials). Data collection took place in a magnetically shielded room using a whole-head MEG system (157 axial gradiometer sensors, 3 reference magnetometers; Kanazawa Institute of Technology, Nonoichi, Japan). Before the experiment, we taped five marker coils on the location of the digitized reference points that help establish the position of the subject’s head before and after the experiment. During the experiment, the participant comfortably lay down in the MEG machine, reading from a screen located approximately 50 cm away with dimmed lights. Text was displayed in a fixed-width Courier New font on a light grey background.

In the experiment, participants were asked to silently read and comprehend short stories consisting of three sentences presented with PsychoPy (Peirce, 2009). The first sentence (context) was displayed as a whole. Participants read this sentence at their own pace and pressed a button to continue. Then a fixation cross (300ms) followed and after a 300ms blank screen the target sentence was presented using Rapid Serial Visual Presentation. Participants were presented with English sentences of 9 words, mostly one word at the time, with the exception of determiner-noun pairs, which were presented together so that the sentence was divided into 7 parts (called ‘words’ from now on). The display time for all words was 300ms. Every word was preceded by a blank screen of 150ms. This was followed by a short third sentence in blue that was either congruent with the previous sentence or incongruent (50%). The continuations were designed such that they targeted the comprehension of different parts of the story (encouraging participants to read the entire narrative with care). One third of the continuations tapped into the modality of the target sentence, in which the continuation is congruent with the modal base (e.g. a sentence about obligation followed by “their mother told them to”) or incongruent with the modal base (e.g. a sentence about obligation followed by “she’s probably right”). We included this manipulation to be sure that participants are paying attention to the fine meaning of the modal target verb. The participant’s task was to press one button with their middle finger for continuations that ‘made sense’ and another button with their index finger if the continuations ‘did not make sense’, after which the next trial started. The participants were instructed to move and blink as little as possible during the task. The trial structure is displayed graphically in Figure 2D.

The experiment consisted of 240 trials in total. The trials were divided into 6 separate blocks (containing 1 item per stimuli set) by a balanced Latin square design and randomized within blocks. Each block consisted of 40 sentences and was presented into two parts during the experiment, resulting into 12 blocks which took about 3-7 minutes each. In between blocks, participants were informed about their overall accuracy. Participants were free to rest in between blocks and were paid $15 (NY) per hour.

#### Data acquisition

MEG data were sampled at 1000 Hz with an online 200 Hz low-pass filter. The signal was offline noise reduced in the software MEG160 (Yokogawa Electric Corporation and Eagle Technology Corporation, Tokyo, Japan) using the signal from the three orthogonally-oriented reference magnetometers (located within the machine, but away from the brain) and the Continuously Adjusted Least-Squares Method (Adachi et al., 2001). Further pre-processing and analysis was performed making use of MNE-Python (Gramfort et al., 2013, 2014) and Eelbrain (Brodbeck, 2017). First, MEG channels that were unresponsive or clearly malfunctioning (separating from all other channels) during the session were interpolated using surrounding channels (6% of the channels in total underwent interpolation, 7-19 channels per participant). We extracted epochs from −2450 to 900 ms relative to the onset of the target verb, which included the entire sentence. The epochs were corrected for the delay between presentation software timing and stimulus presentation, by taking into account the average delay as measured with a photodiode. The data were filtered offline with a band-pass filter between 1 and 40 Hz. Eye blinks and heartbeat artefacts were removed by the use of Independent Component Analysis (ICA) via the “fastICA” option implemented in MNE python (Gramfort et al., 2014). Additionally, we removed a known artefact pattern (‘the iron cross’) that was present at that time across all NY recordings due to an electromagnetic noise source from nearby cables. Any epoch that had a sensor value that was higher than 3pT or lower than −3pT were automatically rejected. Additionally, trials were rejected after visual inspection if multiple channels were affected by obvious noise patterns that exceeded the boundaries of the epoch’s window. In total, this resulted in a trial-rejection rate of 4.6% across the experiment. Baseline correction was performed using data from the 200 ms before the first word of the sentence.

The location of sources was estimated by co-registration of the digitized head shape with the FreeSurfer average brain (Fischl, 2012). A source space containing 2562 sources per hemisphere was constructed for each subject, and a forward solution was created with the Boundary Element Model method. The inverse operator was calculated based on the covariance matrix from the 200 ms pre-stimulus baseline period of the cleaned trials. This inverse operator was applied to the average evoked responses to obtain a time course of minimum norm estimates at each source for each condition (SNR = 3). The direction of the current estimates was freely oriented with respect to the cortical surface, and thus all magnitudes were non-negative. The source estimates were then noise-normalized at each source (Dale et al., 2000), generating dynamic statistical parameter maps (dSPM) that were used in statistical analyses.

#### Statistical Analyses

##### Behavioral data

Responses and reaction times to the 6000 (25×240) congruency decisions were collected and overall accuracy was determined based on the responses to all items. The overall accuracy was used to exclude participants if they scored below 70%. We also examined the accuracy of the 2000 modal task items.

##### MEG data

MEG data were analyzed both with an ROI analysis and with a full-brain analysis, given the explorative nature of our question.

##### ROI Analysis

Since there is no prior neuroimaging work on modal displacement, our ROIs were defined based on previous literature looking at the neural bases of Theory of Mind (Koster-Hale et al., 2017; Mahy et al., 2014; Schurz & Perner, 2015), and included the Inferior Parietal Sulcus (IPS), Temporo-Parietal Junction (rTPJ), Superior Temporal Sulcus (STS), Posterior Cingulate Cortex (PCC), rostral Anterior Cingulate Cortex (rACC) and medial Prefrontal Cortex (mPFC) bilaterally. These functional regions were translated into labels for (bilateral) areas mapped onto the FreeSurfer aparc (Desikan et al., 2006) parcellation (Table 1). Each source current estimate was mapped onto a parcellation, and then averaged over all the sources in each ROI.

**Table 1.**
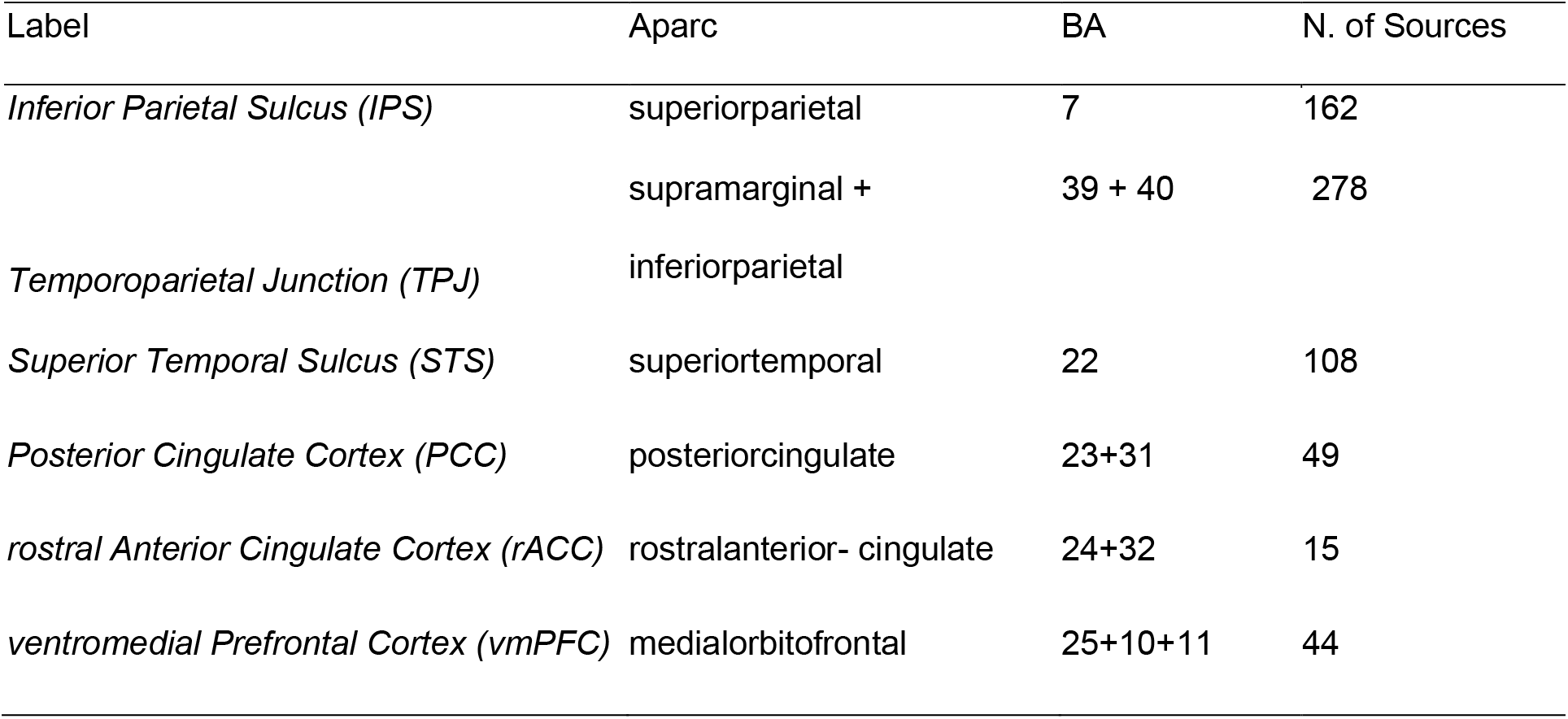
Overview of regions of interest (ROIs) based on the aparc parcellation, with approximately corresponding Brodmann Areas (BA) and number of sources.

The effect of the experimental manipulations on our ROIs was assessed with a cluster-based permutation test (Maris & Oostenveld, 2007), aimed to identify temporal clusters that were affected by our experimental paradigm, corrected for multiple comparisons. We performed a temporal cluster-based permutation mass univariate 2 × 3 repeated-measures analysis of variance (ANOVA) with factors MODAL BASE and FORCE. Since we had no clear predictions about the possible timing of an effect, we used the generous time window of 100-900 milliseconds after the target verb’s onset. Since several trials got rejected during data pre-processing, to ensure comparable SNR across conditions we equalized trial count across conditions (M=36 trials/condition, range=31-39trials/condition)

Our temporal permutation clustering test was performed in Eelbrain 0.27.5 (Brodbeck, 2017) with a standard procedure. An uncorrected ANOVA was fitted at each time point in the analysis time window (100-900 ms). Temporal clusters were formed and chosen for further analysis when *F*-statistics corresponded to significance exceeded the critical alpha-lavel of .05 (uncorrected) for contiguous time points of at least 25 milliseconds. A test statistic corresponding to the cluster magnitude was then determined by summing over all the F-values contained within them then selecting the largest of the cluster-level statistics. Conditions were re-labeled, and test statistics were calculated for each subject for 10,000 times to form a null distribution of the test statistics. The observed clusters were compared to this null distribution and were assigned corrected *p*-values reflecting the proportion of which random partitions resulted in an *F-*statistic greater than the observed *F*-statistic. Since in this method, the time point clusters initially chosen for further analysis are uncorrected, the borders of the clusters should be interpreted as having an approximate nature, not making claims about the *exact* latency or duration of any effects (see Sassenhagen and Draschkow, 2019). Finally, in order to also correct for comparisons across multiple ROIs, we applied a False Discovery Rate correction for multiple comparisons (Benjamini and Hochberg, 1995).

##### Whole Brain Analysis

To complement our ROI analysis, we conducted a full brain analysis, which both described the full spatial extent of any effects observed in the ROI analysis and provided us with information about any effects not captured by the ROI analysis. We performed a spatiotemporal clustering test almost identical to the temporal cluster test described above, only now without averaging sources within an ROI. Instead, an *F*-statistic was calculated for each time point in each source, and spatiotemporal clusters were identified where significance exceeded a *p* value of .05 for at least 10 spatially contiguous sources and for at least 25 milliseconds. Again, following Sassenhagen and Draschkow (2019), the temporal and spatial properties of the identified significant spatio-temporal clusters should be interpreted as an approximate description.

### Experiment 2

#### Participants

The experiment took place on New York University’s New York (NY) and Abu Dhabi (AD) campuses. 24 right-handed, native English speakers participated in the experiment (8 male, 12 AD). Four participants were excluded (1 for not finishing the experiment due to a technical complication, 1 for excessive channel loss and 2 for extreme noise during recording, rendering the data unusable). The age range of the remaining 20 participants was 19-42 years old (M= 26, SD = 6.46). All participants had normal or corrected to normal vision, no history of neurological impairment and provided informed written consent. To mitigate our participant loss, we did not exclude participants based on behavioral accuracy. Participants were pseudo-randomly assigned one of three experimental lists, such that participants were equally divided over each experimental condition.

#### Stimuli

We developed a similar experimental paradigm as Experiment 1, now manipulating the information value of the sentential context rather than manipulating properties of the modal items (modal base and force). We constructed 40 sets of bi-clausal English sentences, containing a causal relationship between the two parts. We contrasted the factual auxiliary verb *do* against the possibility modal verbs *may* and *might*, keeping modal force consistent across items. Sentences differed in their informative content and came in three types: FACTUAL e.g. “Knights carry large swords, so the squires do too”, which introduced novel information with certainty, CONDITIONAL e.g. “If knights carry large swords, the squires do too”, which introduced novel information with uncertainty (indicated by *if*), and PRESUPPOSED, e.g. “Since knights carry large swords, the squires do too”, which introduced presumed to be known information (indicated by *since*) with certainty. The main manipulation (FACTUAL vs CONDITIONAL) was added to test whether a possible effect of belief updating (expected to be present when encountering the factual target verb in the factual condition) disappeared if the information update built on uncertainty (conditional condition). For processing modal displacement, we did not expect a possible effect to be influenced by sentential certainty. We included the PRESUPPOSED condition for exploratory purposes. Each sentence was preceded by a context word, indicating the theme of the upcoming sentence, e.g. “CASTLE”, to stay consistent with Experiment 1, where utterances were preceded by a context sentence. The complete stimulus design and predictions are displayed in Figure 3A. Each stimulus set consisted of 9 sentences (3×3, TYPE: [factual, conditional, presupposed] × VERB: [*may*, *might, do*]) adding to a total of 360 sentences for all 40 stimuli sets (Figure 3B).

**Figure 3.**
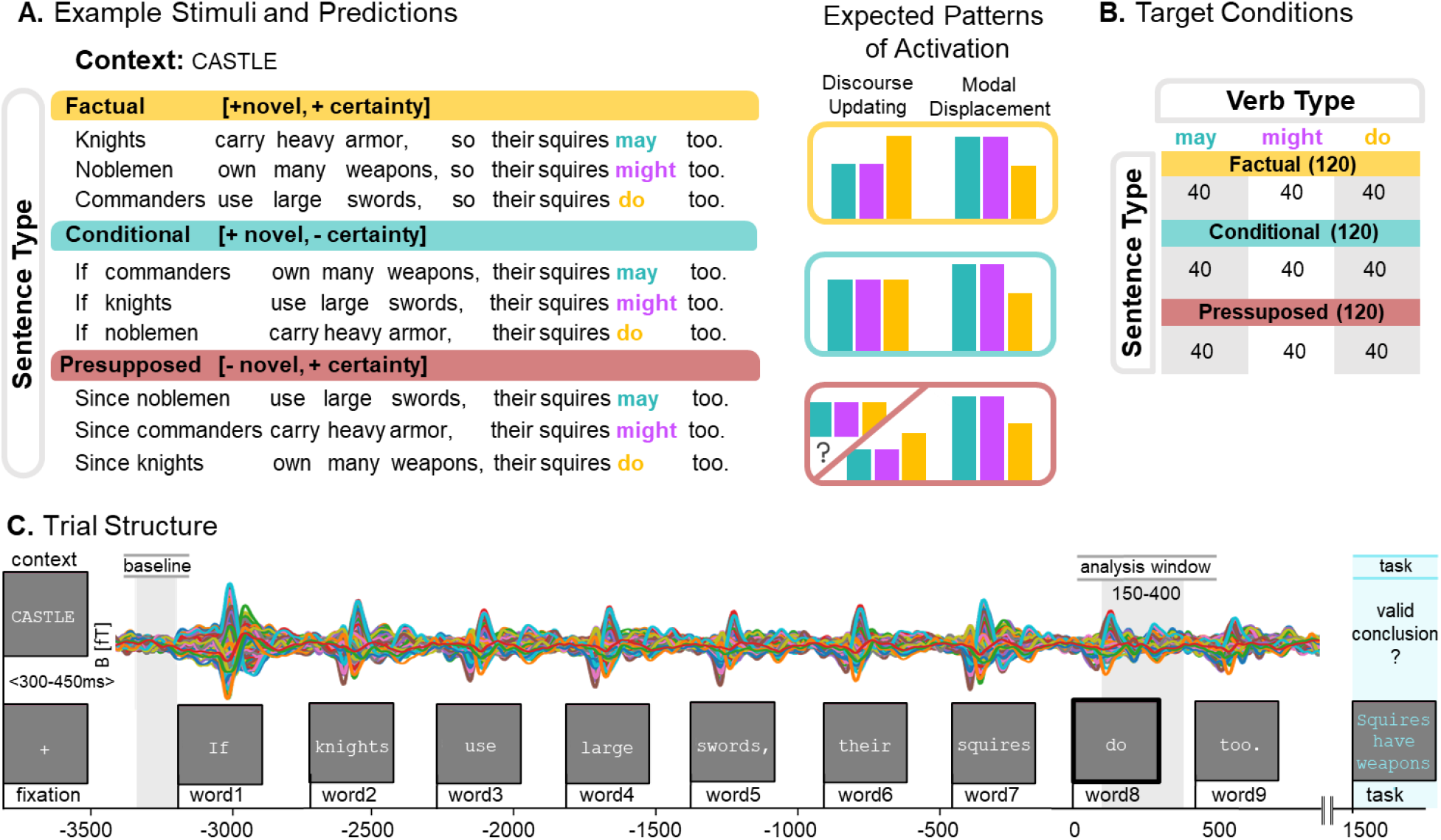
Experimental design and procedure Experiment 2. **A**: Example stimuli set and Predictions. All stimuli were bi-clausal sentences of three different types: factual (*p* so *q*), conditional (if *p* → *q*) and presupposed (since *p* → *q*). These sentence types differed in whether they express information that is novel and certain (factual), novel and uncertain (conditional) or known and certain (presupposed). Each sentence contained either the factual verb *do* or the modal verbs *may* or *might*. Included are expected activation patterns for each verb per sentence type under processes of belief updating and modal displacement. We expect belief updating to take place in factual contexts but not in conditional contexts. For presupposed contexts we had no clear predictions. Activity relate to modal displacement is not expected to change across different sentential environments. **B**: Experimental design with number of items per condition in brackets (total = 360). The stimuli vary among two dimensions: SENTENCE TYPE [factual, conditional and presupposed] and VERB [may, might, do]. **C**: Trial structure with evoked MEG responses from one participant. Procedure similar to Experiment 1. Time windows for baseline correction (−3350 to −3200ms) and statistical analysis (150-400ms) are relative to the target verb (word8) onset.

All utterances were equal in length. Since we pursued a within-participants design and the different sentence conditions within a stimulus set differed minimally, we introduced controlled variance in the first clause of the utterance to make the paradigm seem less repetitive. We constructed three semantically related variants of the subject (e.g. *knights*, *noblemen* and *commanders*) and main event (e.g. *carrying heavy armor*, *owning many weapons* and *using large swords*) that were matched across conditions in a stimulus set so that each subject and action occurred in each of the 9 conditions once. We made three different versions of the experiment such that across versions each condition occurred with all the subject and event variants. Sentential subjects denoted generic groups (e.g. *knights* or *loyal supporters*) and personal/company names (such as *Lisa* or *Facebook*).

#### Procedure

Before recording, we digitized the head shape of each participant with either a FastSCAN laser scanner or a FASTRAK 3D digitizer (Polhemus, VT, USA), following the same procedure as laid out in for Experiment 1. Before participants entered the MEG-room they received verbal instructions and did a short practice block of seven trials. Data collection took place in a magnetically shielded room using whole-head MEG system with 157 (NY) or 208 (AD) channels (Kanazawa Institute of Technology, Kanazawa, Japan). Stimuli were projected onto a screen located above the participant. We made sure to keep the visual angle across both systems consistent, at approximately 0.5° vertically.

In the experiment, participants were asked to silently read and comprehend causally linked sentences presented with PsychoPy (Peirce, 2009), font and background settings identical to Experiment 1. First, a context word was displayed for 600ms followed by a blank screen which display time varied between 300-450 ms. This jitter in display time was included to approximate the temporal variety in Experiment 1 induced by self-paced reading of the context sentence. Then, a fixation cross (300ms) followed and after a 300ms blank screen the target sentence was presented using Rapid Serial Visual Presentation. Participants were presented with English sentences of 9 words, one word at the time (300ms on and 150ms off). This was followed by a conclusion (displayed in blue) that was either a valid conclusion based on prior information (50%) or not. This task was designed such that participants had to pay close attention to the fine details of the target utterances. Forty percent of the questions specifically tapped into the certainty of the prior statement (e.g. the sentence “If knights own many weapons, their squires do too” followed by the valid conclusion “Potentially, the squires own many weapons” or invalid conclusion “The squires own many weapons”). Half of these certainty-based conclusions targeted the first clause of the sentence, while the other half targeted the second half. The other conclusions (60%) were more general e.g. “Knights have (no) squires”. The participant’s task was to press one button with their middle finger for conclusions that were valid and another button with their index finger if the conclusions were invalid, after which the next trial started. The participants were instructed to move and blink as little as possible during the task. The trial structure is displayed in Figure 3C.

The experiment consisted of 360 trials in total. The trials were divided into 9 separate blocks (containing 1 item per stimuli set) using a balanced Latin square design and randomized within blocks. Each block consisted of 40 sentences and was presented in two parts during the experiment, resulting in 18 blocks which took about 3-5 minutes each. In between blocks, participants were informed about their overall accuracy. Participants were free to rest in between blocks and were paid $15 (NY) or 60 AED (AD) per hour.

#### Data acquisition

The same acquisition profile was maintained across both NY and AD systems, with settings as described for Experiment 1. Preprocessing used the same software and pipeline as described for Experiment 1. In total, 7% of the channels were interpolated due to being unresponsive or clearly malfunctioning (NY: 7-14 per participant; AD: 0-18 per participant). We extracted epochs from - 3500 to 1200 ms relative to the onset of the target verb, which included the entire sentence, and rejected epochs containing signal amplitudes that exceeded a threshold of 3 pT (NY) or 2 pT (AD). The NY threshold is higher since that city and system has higher levels of overall ambient magnetic noise. In total, this resulted in a trial-rejection rate of 3.9% across all participants (NY: 5.0%; AD: 2.0%). Baseline correction was performed using data from −3350 to −3200ms relative to the onset of the target verb, before the first word of the sentence. Source estimation followed the exact procedure as described for Experiment 1. The inverse operator was calculated based on the covariance matrix from the 150 ms pre-stimulus baseline period of the cleaned trials.

#### Statistical Analyses

##### Behavioral data

Overall accuracy per participant was based on responses to all 360 items. We also calculated the accuracy of the subset of task items (40%) probing the certainty of the target utterances.

##### MEG data

In order to compare our results from Experiment 1 and 2, we conducted two analyses: an ROI analysis using the regions of interest as defined for Experiment 1 and a conceptual replication analysis searching for spatiotemporal clusters within a predefined region and time window based on the putative discourse updating effect of Experiment 1.

##### ROI Analysis

We used the same ROIs as used for the analysis of Experiment 1, again assessing the effect of our experimental manipulations with a cluster-based permutation test (Maris & Oostenveld, 2007). We performed a temporal cluster-based permutation mass univariate 3 × 3 ANOVA with factors SENTENCE TYPE and VERB. We based our analysis time window on the results of Experiment 1, using a 150-400 ms time window after the target verb’s onset to replicate the effect found in the first experiment. Again, we equalized trial count across conditions. The number of trials per condition that were analyzed was on average 36 out of 40 for NY data (ranging from 31-38 per participant) and 38 out of 40 for the AD data (ranging from 34-40 per participant).

Our temporal permutation clustering test was performed with the same procedure as laid out for Experiment 1 and corrected for comparisons across multiple ROIs (Benjamini & Hochberg, 1995).

##### Conceptual Replication Analysis

With the expectation of replicating the results from Experiment 1, we limited our analysis to the factual sentence type condition. Then, we performed a spatiotemporal clustering analysis using the same procedure and settings as Experiment 1. Informed by the results of Experiment 1, instead of searching through the whole brain, the spatiotemporal analysis was now constrained to a predefined parcellation that combined regions in which we detected the effects of modal force in Experiment 1. This region of interest combined the right banks of superior temporal sulcus and right superior parietal, supramarginal, superior temporal, inferior parietal and middle temporal gyri from the Freesurfer aparc parcellation. Like the ROI analysis, the time window of interest was 150-400 ms after the verb’s onset.

## RESULTS

### Experiment 1

#### Behavioral Results

The mean overall accuracy for the story congruency task was 83.1% (SD = .05), ranging from 71.6%-92.5% across participants. The accuracy of the one third of the congruency task items that tapped into modality was 73.3% (SD = .08) ranging from 60.0 - 88.8% across participants, and was substantially lower than the accuracy of the other general items, which was 87.9% (SD = .05) ranging from 74.4 - 94.4% across participants.

#### ROI Results

We ran a 2 (MODAL BASE: knowledge-based, rule-based) by 3 (MODAL FORCE: possibility, necessity, factual) within-subjects temporal ANOVA for the ROIs specified for Experiment 1. Since *may* and *must* differ in their lexical frequency across modal bases (*may* is high frequency as knowledge-based modal and low frequency as rule-based modal, *must* low frequency as knowledge-based modal and high frequency as rule-based modal, see ‘Stimuli’) we only report results that show consistent results across the *force* manipulation (knowledge-based and rule-based *may* or *must* patterning together) or the *modal base* manipulation (*may* and *must* patterning together).

The ANOVA revealed a significant effect of modal force in the right Inferior Parietal Sulcus (rIPS) within our test window of 100-900 ms after the target verb’s onset (*p* = .046), where the factual condition (*do*) elicited more activation than the modal (*may* and *must*) conditions. This temporal cluster extended from approximately 280-340 ms. We observed a similar effect in a temporal cluster in the right Temporo-parietal Junction (rTPJ) around 240-275 ms, although this effect only survived multiple comparisons correction across time, not across multiple regions of interest (uncorrected p = .054, p = .13). Additionally, we found a trending effect of modal force in the right rostral Anterior Cingulate Cortex (rACC), with increased activation for the necessity modal *must* over the other conditions (uncorrected *p* = .008, *p* = .099). We did not observe any other clusters in the remaining ROIs of the right hemisphere and did not observe any clusters in the left hemisphere. We summarized the ROI results in Figure 4 by depicting the activation patterns of the detected reliable clusters. The measured activity for each of the ROIs over our time window of interest are displayed in Figure 5.

**Figure 4.**
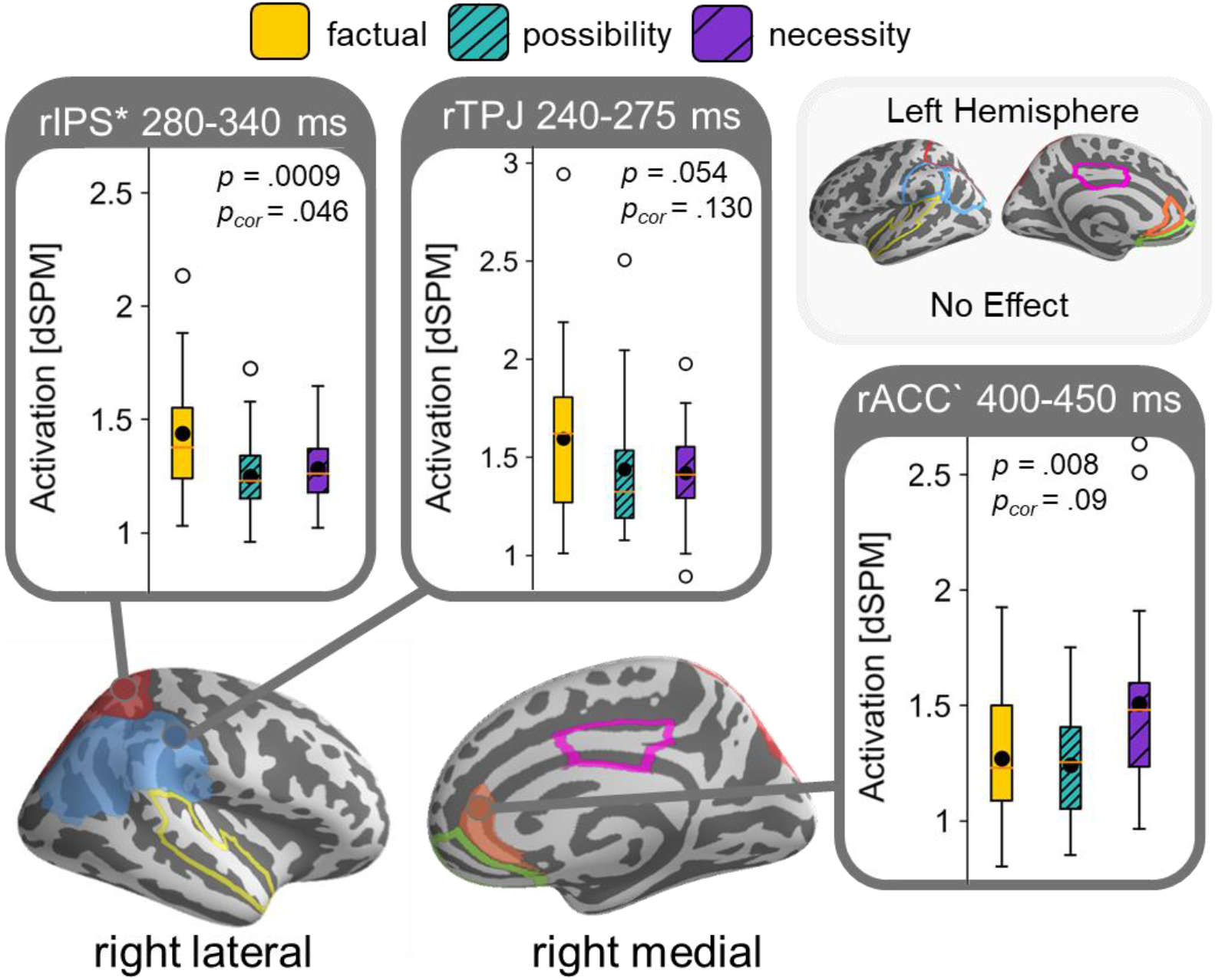
Summary Region of Interest (ROI) Results Experiment 1 showing a main effect for factual over modal conditions in right IPS and TPJ, and an increase in activation for necessity in the rACC. Results collapsed for MODAL BASE (knowledge-based and rule-based modals grouped together). Boxplots of estimated brain activity within the time window of the identified temporal clusters, black dots indicate mean activity. Regions of interest outlined on brain and shaded when containing identified clusters. Clusters significant after correction comparison across multiple ROIs indicated with asterisk and with grave accent when trending.

**Figure 5.**
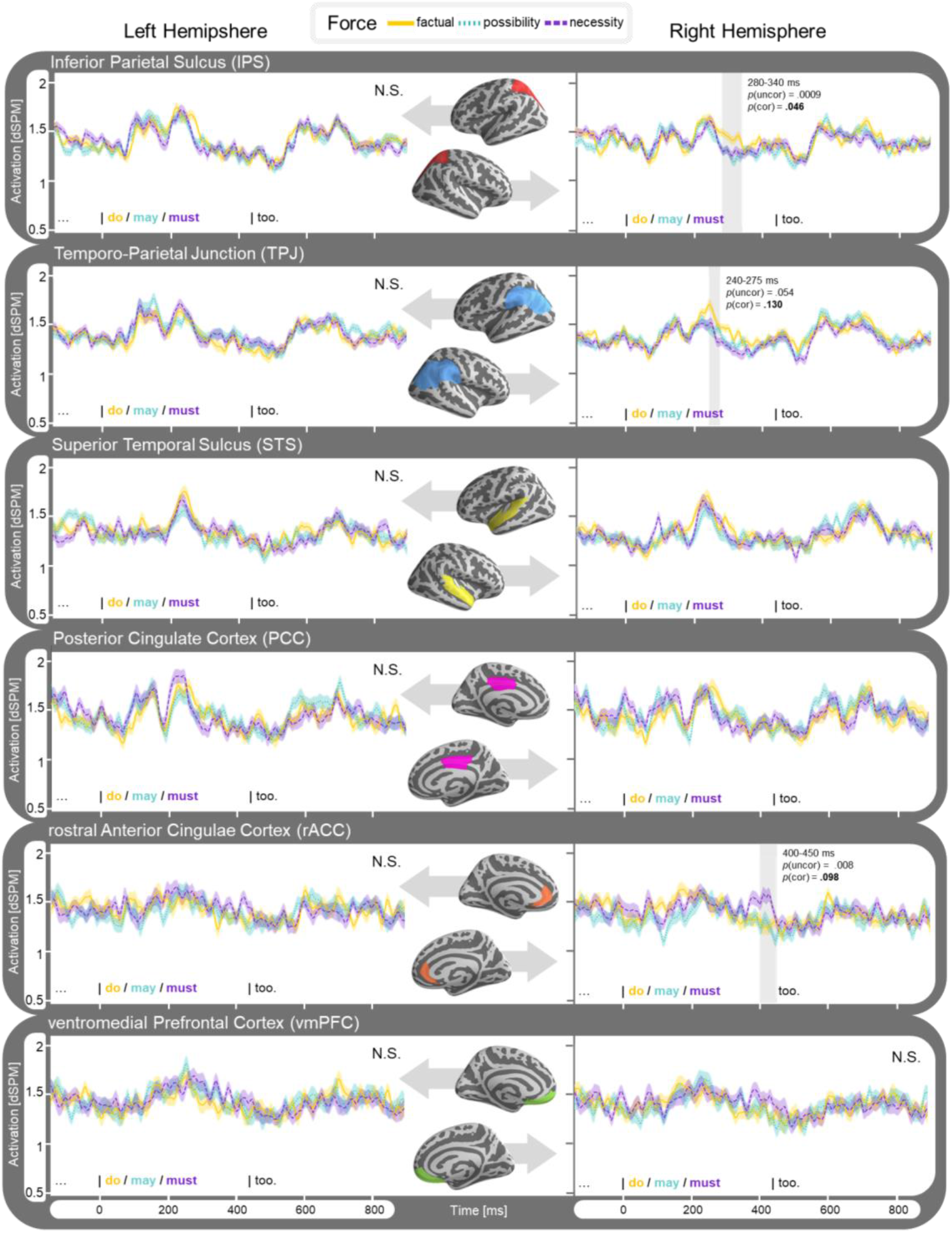
Time course of estimated average activity [dSPM] per ROI of Experiment 1. Left hemisphere ROIs displayed on the left side, and right hemisphere on the right. Results collapsed for MODAL BASE (knowledge-based and rule-based modals grouped together). Detected clusters within time window 100-900ms highlighted and significance indicated for effect within cluster (p_uncor_) and when corrected for comparison across multiple regions (*p*_cor_).

#### Spatiotemporal Results (Whole Brain)

A full-brain analysis revealed a significant effect for modal force, eliciting stronger activity for our factual condition over our modal conditions (p= .033) in our 100-900 ms time window. We detected a cluster between approximately 210-350 ms centering around the right Temporoparietal Junction (rTPJ) extending posteriorly over to the right Intraparietal Sulcus (rIPS) to the medial cortex, covering the cuneus, parts of the precuneus, and ending in the posterior cingulate cortex (Figure 6). The activation in this cluster reflects the activity we found for the effect of modal force in the rIPS and rTPJ of our ROI-analysis. No other significant clusters were found.

**Figure 6.**
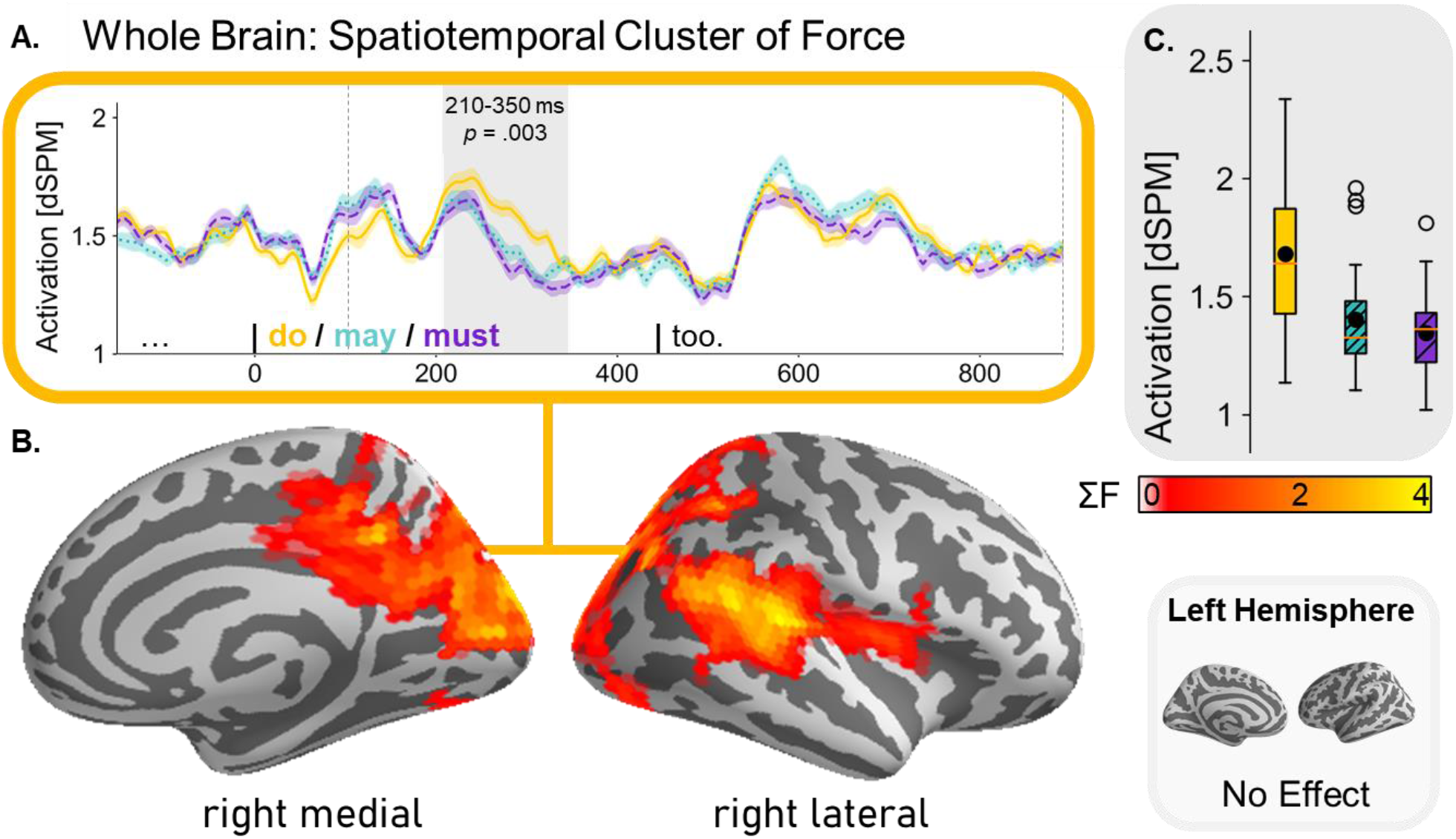
Identified spatiotemporal cluster of whole-brain analysis Experiment 1. **A.** Time course estimated brain activity [dSPM] and identified cluster (in grey). Boundaries analysis window (100-900 ms) indicated by dashed lines. **B.** FreeSurfer average brain shows spatial distribution of cluster, color shading indicating the sum of cluster-level F statistic (gained from cluster-based permutation test). **C.** Boxplots of estimated brain activity (factual > modal) within the identified time window of the spatiotemporal cluster, black dots indicate mean activity.

### Experiment 2

#### Behavioral Results

The mean overall accuracy for the conclusion validation task was 85.6% (SD=.09), ranging from 64.7%-96.9% across participants. The accuracy of the subset of the validation task items that tapped into certainty was 83.7% (SD = .10) ranging from 57.6 - 95.2% across participants.

#### ROI Results

We ran a 3 (SENTENCE TYPE: factual, conditional, presupposed) by 3 (VERB: may, might, do) within-subjects temporal ANOVA for the same ROIs specified for Experiment 1. We only observed effects that survived multiple comparisons correction across time, but not across multiple regions of interest. The ANOVA revealed an interaction effect of VERB and SENTENCE TYPE in the left anterior Cingulate Cortex (lACC) within our test window of 150-400 ms after the target verb’s onset (uncorrected p = .034, p = .341), where the factual condition (*do*) elicited more activation than the modal (*may* and *must*) conditions in factual sentences, but not in conditional or presupposed sentences. In fact, in presupposed sentences the factual condition elicited less activity than the modal conditions. The temporal cluster reflecting this activity difference extended from approximately 365-395 ms. We observed a similar effect in a temporal cluster in the right ventromedial Prefrontal Cortex (rvMPFC) around 345-370 ms (uncorrected p = .032, p = .327). No other clusters were detected in any of the other regions of interest. We summarized the ROI results in Figure 7 by depicting the time course of the detected reliable clusters. The effect in the lACC was most prominent in the NY data while the effect in the rvMPFC was more prominent in the AD data (Appendix II). The measured activity for each of the ROIs over our time window of interest in the factual sentential context (for comparison with Figure 5) is displayed in Figure 8.

**Figure 7.**
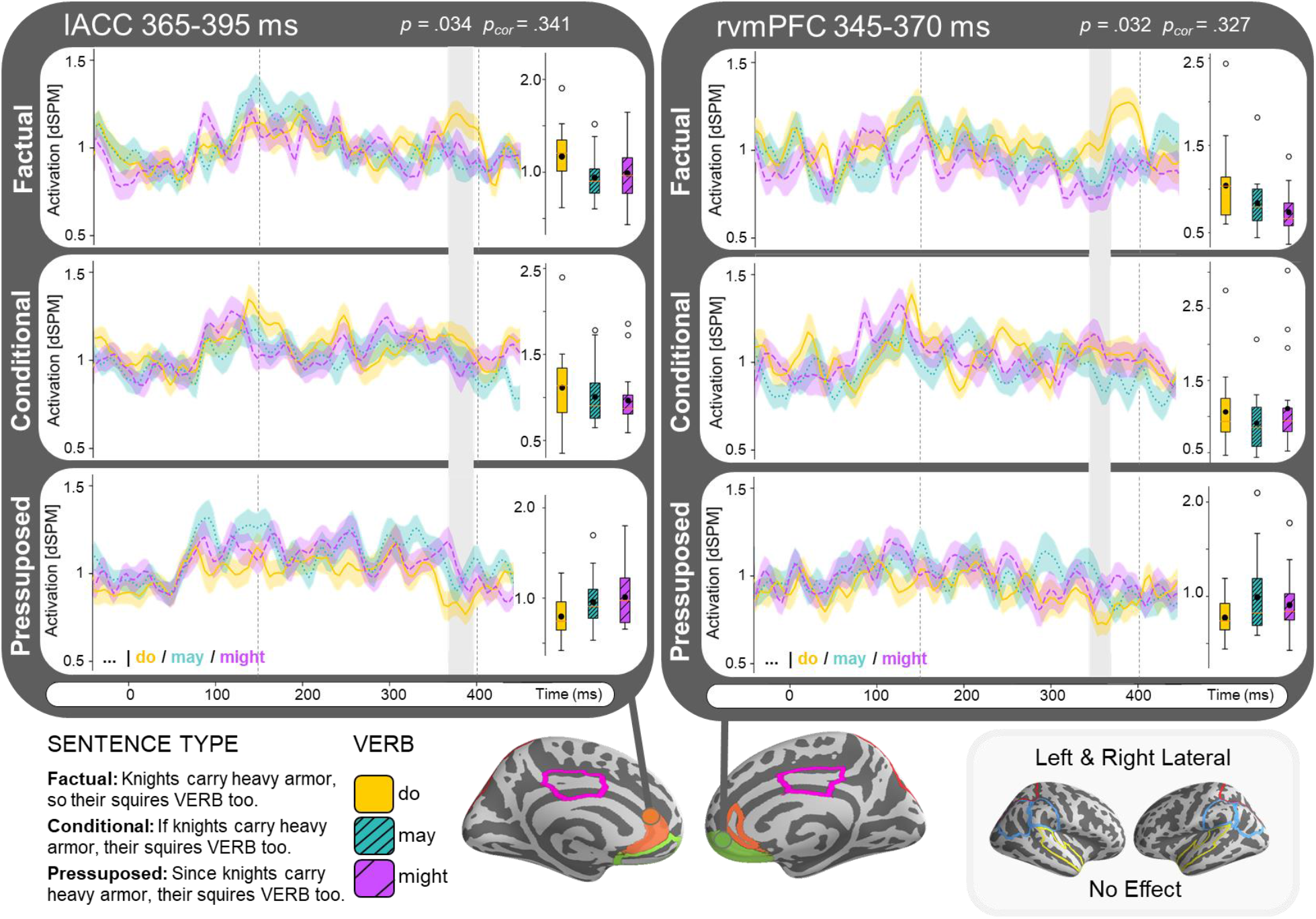
Time course estimated brain activity [dSPM] of reliable detected clusters from ROI analysis Experiment 2. Both the lACC and rvmPFC show an interaction between sentence type (factual, conditional and presupposed) and verb (*do*, *may* or *might*) with increased activation for *do* > *may/might* when embedded in factual sentences, and decreased activation for *do* < *may/might* in presupposed sentences. Boundaries of the analysis window (150-400 ms) are indicated by dashed lines, identified clusters displayed in grey. Boxplots of estimated brain activity within the time window of the identified temporal clusters, black dots indicate mean activity. Regions of interest outlined on brain and shaded when containing identified clusters. Cluster effects not significant after correction comparison across multiple regions of interest. The effect in the lACC was most prominent in the NY data while the effect in the rvMPFC was more prominent in the AD data (Appendix II).

**Figure 8.**
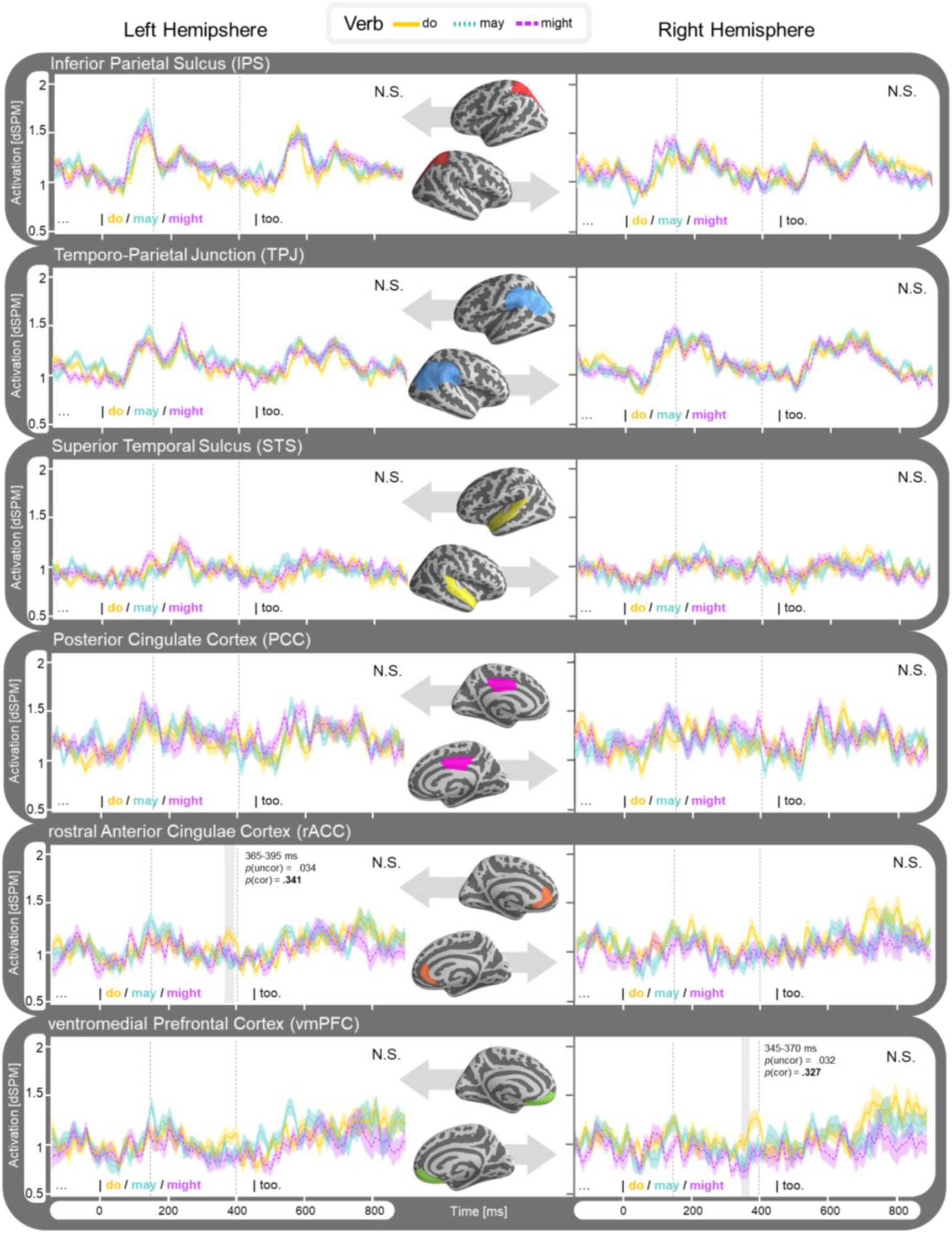
Time course of estimated average activity [dSPM] per ROI of Experiment 2 for factual sentence type (*p* so *q*). Left hemisphere ROIs displayed on the left side, and right hemisphere on the right. Results collapsed for MODAL BASE (knowledge-based and rule-based modals grouped together). Detected clusters within time window 150-400ms (indicated with dashed lines) highlighted and significance indicated for effect within cluster (p_uncor_) and when corrected for comparison across multiple regions (*p*_cor_).

#### Conceptual Replication Results

We performed a spatiotemporal clustering test in the time window 150-400 ms in a region of interest covering right lateral temporoparietal areas aiming to replicate the effect found in Experiment 1. Unlike the results of Experiment 1, a one-way ANOVA comparing activity within the VERB condition (*do*, *may* and *might*) in FACTUAL sentences detected no significant clusters in this area. This corroborates the results of the ROI analysis, in which we similarly found no difference in activity between the factual and modal verbs in the right IPS, TPJ or STS.

## DISCUSSION

In this work, we conducted two experiments to explore the neural correlates of modal displacement and discourse model updating during language comprehension. During natural discourse comprehension, the comprehender does not only integrate incoming factual information into an evolving discourse model, but also entertains hypothetical situations denoted with modal utterances. We investigated how the brain distinguishes between factual and modal information.

Our stimuli contained short scenarios with two parts. The first part of the narrative established some property or habit that applied to one entity (e.g. “Knights carry heavy armor”), The second provided additional information about a second entity that was either factual (e.g. “the squires *do* too”) or modal (e.g. “the squires *may*/*must*/*might* too”). While the factual utterances indicated an actual change in situation, requiring the discourse representation to be updated, the modal utterances merely indicated a possible (uncertain) change. Our data showed that the factual condition elicited reliably stronger activation than the modal condition in right temporoparietal (Experiment 1) and medial frontal regions (Experiment 2). Below we discuss these increases as possible neural correlates of discourse model updating, elicited in the presence of updates that are certain (factual) but not for updates that are uncertain (modal).

### Neural Correlates of Discourse Updating

Discourse updating, the operation of updating the mental representation of a situation, was modelled here as the attribution of a property to a new entity. Prior behavioral research has shown that mental representations of discourse are dynamically updated when presented with new facts (Glenberg et al., 1987; Morrow et al., 1989; Zwaan & Madden, 2004). Such modal updating has been associated with increased activation in the mPFC, PCC and temporo-parietal areas (Ferstl et al., 2005; Fletcher et al., 1995; Speer et al., 2007; Xu et al., 2005; Yarkoni et al., 2008). In Experiment 1, we found an increase in source-localized MEG responses for factual over modal statements. Specifically, activity increased in factual statements in the right lateral temporal and parietal hemisphere at approximately 200-350 ms after target verb onset. This effect was most pronounced in the right inferior parietal sulcus (rIPS) and less so in the right temporo-parietal junction (rTPJ). This pattern of activity is compatible with behavioral findings on discourse updating. Factual utterances signal an actual change in the discourse, and when this information is incorporated into the comprehender’s mental representation this results in increased brain activity. In contrast, modal utterances only indicate a possible change of situation. Since the update is uncertain, situation model updating does not take place.

In Experiment 2, we manipulated the broader sentential context in which novel factual and modal information was presented. In contrast to Experiment 1, where the target sentence always built on a certain factual base, we now also presented the target utterance in conditionals that were hypothetical (uncertain, i.e. ‘If knights carry large swords…) or presupposed (presumed to be common knowledge, i.e. ‘Since knights carry large swords…). We expected discourse updating to only take place when the situational change is certain, and that embedding a factual update into a hypothetical conditional should prevent discourse updating from taking place due to the entire scenario being uncertain (Figure 1).

While Experiment 2 was designed to replicate the results from Experiment 1 with our factual sentential context, we instead found that this time our ROI analysis (using the same regions of interest as defined for Experiment 1) revealed no differences in activity between factual and modal utterances in the right lateral hemisphere. This was confirmed by a replication analysis searching for spatiotemporal clusters targeting right lateral temporoparietal areas within the time window of 150-400 ms. Instead, we now found increased activity for factual over modal conditions in a temporal cluster in two adjacent areas: the left anterior Cingulate Cortex (lACC) and right ventromedial Prefrontal Cortex (rvmPFC) within our test window of 150-400 ms after the target verb’s onset. This effect only survived multiple comparisons correction across time, not across multiple regions of interest. The hypothesis that this activation reflects discourse updating gains weight from the fact that we only observed this pattern of activity when the sentential context was factual (“Knights carry large swords, so their squires *do*/*may*/*might* too.”) but not when the sentential context was hypothetical (“If knights carry large swords, their squires *do*/*may*/*might* too.”). This would be in line with the idea that discourse model updating only takes place under certain situational changes, though such a conclusion has to be drawn with caution, as the results of Experiment 2 were not that robust.

This presumed discourse updating effect resonates with prior behavioral studies on discourse updating and situation model maintenance. Discourse models representing a situation are dynamically updated as novel information indicating a change of situation comes along. As a consequence of model updating, ‘old’ information that is no longer relevant to the *here-and-now* of a story is backgrounded, which is measurable in longer retrieval times in probe-recognition tasks compared to information that is still relevant to the current situation (Glenberg et al., 1987; Morrow et al., 1989; Zwaan & Madden, 2004). De Vega et al. (2012; 2007) investigated whether this model updating also takes place when integrating hypothetical information, comparing accessibility after encountering factual (“As he had enough time, he went to the café to drink a beer”) and counterfactual utterance (“If he had enough time, he would have gone to the café to drink a beer”). De Vega et al. (2007) found evidence for discourse updating when integrating factual information but not for counterfactual information, leading them to conclude that the hypothetical meaning of counterfactuals does not contribute to the build-up of the discourse representation. This finding was corroborated in an ERP study, where increased negativity after factual compared to counterfactual continuation utterances and reduced gamma power following counterfactuals were taken to indicate that the counterfactual’s ‘as if’ meaning is not integrated into the discourse (de Vega & Urrutia, 2012). Our results likewise suggest that mental model updating takes place for the integration of novel factual information, but not for hypothetical information as indicated by modality (*may/must/might*) or conditionality (*if…*).

While the results of Experiment 2 are less strong, they address some possible alternative explanations for the robust effect observed in Experiment 1, which we hypothesized to reflect discourse updating. One might wonder whether a more low-level explanation could explain the observed activity increase for *do* over *may* and *must* in the first experiment, such as an inherent difference in lexical frequency (*do* is more frequent than *may* and *must*), polysemy (*may* and *must* are polysemous while *do* is not) or type of ellipsis (*do* ellipsis syntax may differ slightly from *may*/*must*). These alternative explanations are contradicted by the results of Experiment 2, as we would have expected low-level effects like these to have been replicated in the same location and be insensitive to the experimental manipulation of our sentential context. Furthermore, we included the non-polysemous modal *might* to rule out the polysemy hypothesis. If the increase of factual over modal conditions in both experiments reflects discourse updating however, the question arises what caused the shift in location of this effect between experiments.

### Updating the Representation of Someone Else’s Mental State versus One’s Own

In both of our experiments, we observed an increase for factual over modal expressions – henceforth “updating effect” – but the effect localized differently across the two experiments. In Experiment 1, the updating effect was found in the rIPS and the adjacent rTPJ, while in Experiment 2 we did not observe any effects in these specific areas. Instead, Experiment 2 elicited a similar pattern of activity in medial frontal areas: the lACC and rvmPFC. Both frontal medial and temporal parietal areas have been found to be involved in constructing and maintaining discourse representations in fMRI studies (Ezzyat & Davachi, 2011; Friese et al., 2008; Speer et al., 2007; Xu et al., 2005; Yarkoni et al., 2008). For example, Xu et al., (2005) investigated natural language comprehension at the level of words, sentences and narratives. When comparing visually presented isolated sentences and narratives, they observed robust response increases in several bilateral brain regions including the precuneus, medial prefrontal and dorsal temporo-parieto-occipital cortices. In a similar manipulation, contrasting unrelated sentences with coherent narratives, Yarkoni et al. (2008) found narrative-specific activation in the mPFC and additional neural contributions of posterior parietal regions supporting situation model construction and frontotemporal regions supporting situation model maintenance.

While both temporoparietal and frontal medial areas are part of the network engaged during narrative comprehension, one may wonder why Experiment 2 did not replicate the discourse updating effect of Experiment 1 in the same regions. The reason for this may be related to a change in materials between the experiments, altering whose mental representation is updated. In Experiment 1, all target beliefs are attributed to a third person character, e.g. “But the king learns that the squires do too”. This third person character was included to enhance the contrast between the knowledge-based and rule-based modal readings, varying between authority and observer figures respectively. In contrast, Experiment 2 lacked this third person character and embedding verb (“…, so the squires do too”) for the target manipulation to appear in conditional structures. By making this change in stimuli, we inadvertently changed whose mental state is updated during comprehension, someone else’s (Experiment 1) or the participant’s own (Experiment 2). As laid out in the Introduction, Theory of Mind encompasses the ability to represent someone else’s mental state and keep this separate from our own. Theory of Mind reasoning engages a network of brain regions, but it has been argued that particularly the right TPJ is involved in representing the mental state of others (Saxe & Kanwisher, 2003; Saxe & Powell, 2006; Saxe & Wexler, 2005; Vistoli et al., 2011) or reorienting attention (Corbetta et al., 2008; Decety & Lamm, 2007; Mitchell, 2008; Rothmayr et al., 2011). We tentatively suggest that the discourse updating effect in Experiment 1 localized around the right TPJ because it involved updating a discourse representation separate from the comprehender’s own. Experiment 2 involved updating one’s own global representation and elicited activation in frontal medial regions. This is in line with studies finding medial prefrontal activity for tasks that require people to reflect on or introspect about their own mental states (Gusnard et al., 2001; Mitchell et al., 2005; Zhu et al., 2007). And is also compatible with Ezzyat and Davachi (2011), who found that the bilateral vmPFC seemed especially engaged when integrating information within events, suggesting that this region could be sensitive to discourse updating.

Alternatively, it could be the case that the difference in results between Experiment 1 and Experiment 2 has to do with the narrative complexity. In Experiment 1, the target sentence appeared after an initial context sentence that was read at the participant’s own pace. In the context sentence, a property was established for one entity. The target utterance then indicated that this property was also (possibly) shared by a second entity. In Experiment 2, the context before the target utterance merely consisted of one word introducing the general setting of the following utterance. The target utterance then consisted of two clauses, the first one establishing a (possible) property for one entity, while the second one stated that this property was (possibly) shared by a second entity. The entire target utterance was displayed with rapid serial visual presentation. Compared to Experiment 1, Experiment 2 thus allowed less time for participants to appreciate the initial situation (property being attributed to one entity) before updating this information (property also being attributed to second entity). Under this alternative account, temporal parietal areas would be more involved with constructing a larger discourse representation (coherence between sentences), while the medial frontal areas would be more involved with initializing a discourse representation. This would be in line with Xu et al. (2005), who observed increased activity in the right hemisphere as contextual complexity increased.

An argument against this alternative hypothesis comes from recent work by Jacoby and Fedorenko (2018) investigating the neural correlates of expository discourse comprehension. While prior studies detected right temporal parietal engagement in comprehension of narratives (stories built around characters), expository texts (constituting facts about the real world) elicited no effect of discourse coherency in posterior ToM regions like the rTPJ (Jacoby & Fedorenko, 2018). This suggests that these regions only engage in coherence building for discourse in which you take someone else’s perspective. However, Jacoby and Fedorenko (2018) did find that the mPFC was sensitive to discourse coherency of expository texts. Since their expository texts were as complex as a narrative, it cannot be the case that the lack of engagement of the rTPJ observed for expository texts is due to a lack of discourse complexity. At the same time, the finding that the mPFC is sensitive to the coherence of expository texts suggests it could be involved in updating one’s own discourse beliefs.

### Neural Correlates of Modal Displacement?

We found no reliable increases in neural activity when modal displacement occurred. However, the fact that we did find neural activation dissociating between the factual and modal condition suggests that participants processed the modal items as being different from the factual ones. Given that the increase in activation of factual over modal conditions takes place during the discourse integration of information indicating an actual change in situation, but not when integrating information regarding an uncertain (hypothetical) change, the most likely interpretation of our data is that this difference in activation reflects discourse updating.

However, if non-factual information does not get integrated into an existing situation model, the question remains how we *do* represent this information. The theoretical background for the current study was that modal displacement would involve the generation of multiple possibilities (von Fintel, 2006; Iatridou, 2000; Johnson-Laird, 1994; Kratzer, 2012). Intuitively, this would suggest that when presented with uncertainty, the comprehender postulates multiple mental representations of these different possibilities, the minimal one being a negated version (if squires *might* sit at round tables, this introduces the alternative possibility that maybe they do not). Considering multiple possibilities in parallel is thought to be cognitively demanding (Leahy & Carey, 2019), and we thus expected additional activity related to this operation. It is possible that this assumption was wrong, and that for example, the decreased activity for modal utterances compared to factual utterance is indicative of modal displacement rather than discourse updating. However, it is difficult to gauge why this modal displacement is dependent on the sentential context and why we would find this correlate shifting in location across experiments. Alternatively, there might not be any correlates of representing multiple possibilities in the cortex at the level we investigated in this paper. Recently, Kay et al. (2020) found that possibility generation in rats involves a constant cycling between possible future scenarios in hippocampal neuron populations. At a constant cycling of 8 Hz the cells alternated between encoding two different possible futures. The authors suggest this finding might extend to the representation of hypothetical possibilities in human brains, possibly extending to brain regions connected to the hippocampus.

Lastly, some have proposed that the representation of modality involves marking a representation with a symbolic operator, indicating that this representation can be neither ruled out nor added into the actual model (Leahy & Carey, 2019). This theory would not require people to actively postulate alternative situations, though the question remains how this uncertain information would be maintained and linked to the prior discourse if not incorporated into the existing situation model. For now, these questions are still open to future exploration.

### No Effect of Modal Base and Force

Our stimuli in Experiment 1 were carefully designed to investigate the online comprehension of modal verbs varying in modal base (knowledge-based versus rule-based) and force (possibility versus necessity). However, we found no reliable effects of these manipulations. We did find an effect in the right Anterior Cingulate Cortex showing increased activation for necessity modals over the other conditions (Figure 5), but this effect only survived multiple comparisons correction across time, not across multiple regions of interest. The rostral ACC is, besides its involvement in ToM tasks, also argued to be involved in error processing and conflict resolution (Dreher & Grafman, 2003; Kiehl et al., 2000), suggesting that our effect may reflect some unnaturalness in our stimuli. The verb *must* requires strong evidence, but the surrounding context was made to be also compatible with weaker evidence (to allow for the appearance of *may*). Possibly, our stimuli contained too little evidence to naturally say *must*, eliciting increased activation in the rACC when resolving this conflict.

## CONCLUSION

This work investigated the integration of factual and modal information into short narratives. While the factual utterances indicated an actual change in situation, requiring the discourse representation to be updated, the modal utterances merely indicated a possible (uncertain) change as these utterances displaced from the narrative’s here-and-now. In a controlled within-subjects design, we measured source-localized MEG responses while participants integrated modal and factual information into a short narrative. While we did not find any regions of the brain more engaged by the modal conditions over the factual conditions (which could reflect engagement with modal displacement), we did find the opposite pattern of activation where certain brain regions elicited stronger activation for the factual over the modal condition. This increase in activation may be a neural correlate of mental discourse representation updating. This activity difference seems to go away as soon as the factual update is presented in an uncertain (conditional) sentential environment, supporting the idea that discourse updating only takes place when the change in the situation is certain. To our knowledge, this was the first attempt to explore the neural bases of displacement. While we have established possible neural correlates of fact comprehension, the question of how uncertain information is integrated into a discourse representation remains open. We hope that our work establishes a starting point for further investigations of this phenomenon.

## Appendix I.

Details on controlled between-stimuli variation.

|  |  |  |  |  |  |  |  |  |  |
| --- | --- | --- | --- | --- | --- | --- | --- | --- | --- |
| e.g. | A<br>CON | the | B<br>NOUN.SG | E<br>VERB1 | that | C<br>DET | NOUN.PL \| **TARGET** | D<br><ELIDED VP> | too |
|  | "But | the | king | says | that | their | squires \| **may** | <sit around the round table> | too." |

**A. Variation of CONNECTIVES across flavors**
| Base | And | But | So | Total |
| --- | --- | --- | --- | --- |
| *factual* | 30 | 28 | 22 | 80 |
| *rules* | 30 | 28 | 22 | 80 |
| *knowledge* | 30 | 28 | 22 | 80 |
| Total | 90 | 84 | 66 | 240 |

**C. Variation of DETERMINERS across flavors**
| Flavor | LD<sup>i</sup> | SD<sup>ii</sup> | the | Total |
| --- | --- | --- | --- | --- |
| *factual* | 22 | 22 | 36 | 80 |
| *rules* | 20 | 24 | 36 | 80 |
| *knowledge* | 24 | 20 | 36 | 80 |
| Total | 66 | 66 | 108 | 240 |

<sup>i</sup>LD indicates long distance pronouns, pronouns that refer back to a referent in the context sentence, e.g. 'Apparently knights overhear a lot of secrets in the castle. But the servant concludes that **their** squires must too'.
<sup>ii</sup>SD indicates a short distance pronoun, pronouns that refer back to a referent in the target sentence, e.g. 'The bride's father is known to give boring speeches at receptions. But the bride hints that **her** brothers may too'.

**B. Variation of NOUNS across modal base conditions**
| Values | Rules | Knowledge |
| --- | --- | --- |
| Average Length (letters) | 6,8 | 7,6 |
| Average Freq_HAL | 83,9 | 87,20 |
| Average Log_Freq_HAL | 8984,9 | 8585,2 |
| Average NSyll | 2,25 | 2,5 |
| Average NMorph | 1,6 | 1,75 |

**D. Variation of ELIDED VP across modal base conditions (Av. = average)**
| Base | Av. Ellipsis<br>Length (words) | Av. Ellipsis<br>Length (letters) | Stative<br>Verbs<sup>i</sup> | Transitive<br>Verbs<sup>ii</sup> | Av. Node<br>Count<sup>iii</sup> | Av. Node<br>Complexity<sup>iv</sup> |
| --- | --- | --- | --- | --- | --- | --- |
| *rules* | 4,975 | 28,15 | 8% | 88% | 2,7 | 2,2 |
| *knowledge* | 4,775 | 27,9 | 15% | 88% | 2,925 | 2,55 |

<sup>i</sup>Percentage of stative verbs (e.g. *know*, *accept*, *live*).
<sup>ii</sup>Percentage of transitive verbs (with two arguments), 12 percent are intransitive verbs (taking one argument).
<sup>iii</sup>Estimation of size syntactic structure by counting structural phrase nodes. Counted nodes are: noun phrase (NP), verb phrase (VP), predicate adjective phrase (AP), preposition phrase (PP), infinitival phrase (IP). NPs within APs or PPs were not counted. IPs were only counted in presence of the infinitival marker 'to'. Adverbs, particles and attributive adjectives were ignored.
<sup>iv</sup>Layers of embedded structure (complements). PPs are considered adjuncts not adding to complexity: e.g. [IP [VP [NP]]] = counted as having 3 structural layers, [IP [VP] [PP]] = counted as having 2 structural layers.

**E1.** Variety and repetition of VERB1 per condition.
| Factual<sup>i</sup> |  | Knowledge<sup>ii</sup> |  | Rules<sup>iii</sup> |  |
| --- | --- | --- | --- | --- | --- |
| acknowledges | 7 | assumes | 7 | agrees | 7 |
| discovers | 7 | believes | 7 | announces | 7 |
| hears | 7 | concludes | 7 | argues | 7 |
| knows | 7 | fears | 7 | assures | 6 |
| learns | 7 | figures | 6 | decides | 7 |
| notices | 7 | gathers | 5 | declares | 7 |
| observes | 7 | hints | 6 | determines | 7 |
| realizes | 6 | presumes | 7 | indicates | 7 |
| recognizes | 7 | reckons | 7 | informs | 7 |
| regrets | 5 | suspects | 7 | says | 7 |
| remembers | 6 | thinks | 7 | states | 7 |
| reveals | 7 | worries | 7 | warns | 4 |
| Total | 80 |  | 80 |  | 80 |

<sup>i</sup> Verbs of saying and directives/desideratives.
<sup>ii</sup> Mental/emotion verbs.
<sup>iii</sup> Perception/(semi)-factive verbs.

**E2.** Variation of VERB1 per modal base condition.
| Values | Factual | Rules | Knowledge |
| --- | --- | --- | --- |
| *Average Length* (letters) | 7.75 | 6,75 | 7 |
| *Average Freq_HAL* | 7886.9 | 8624.2 | 8016.9 |
| *Average Log_Freq_HAL* | 141.6 | 111.53 | 111.598 |
| *Average NSyll* | 2.417 | 1.912 | 1.75 |
| *Average NMorph* | 2.25 | 2.33 | 2.25 |

## Appendix II

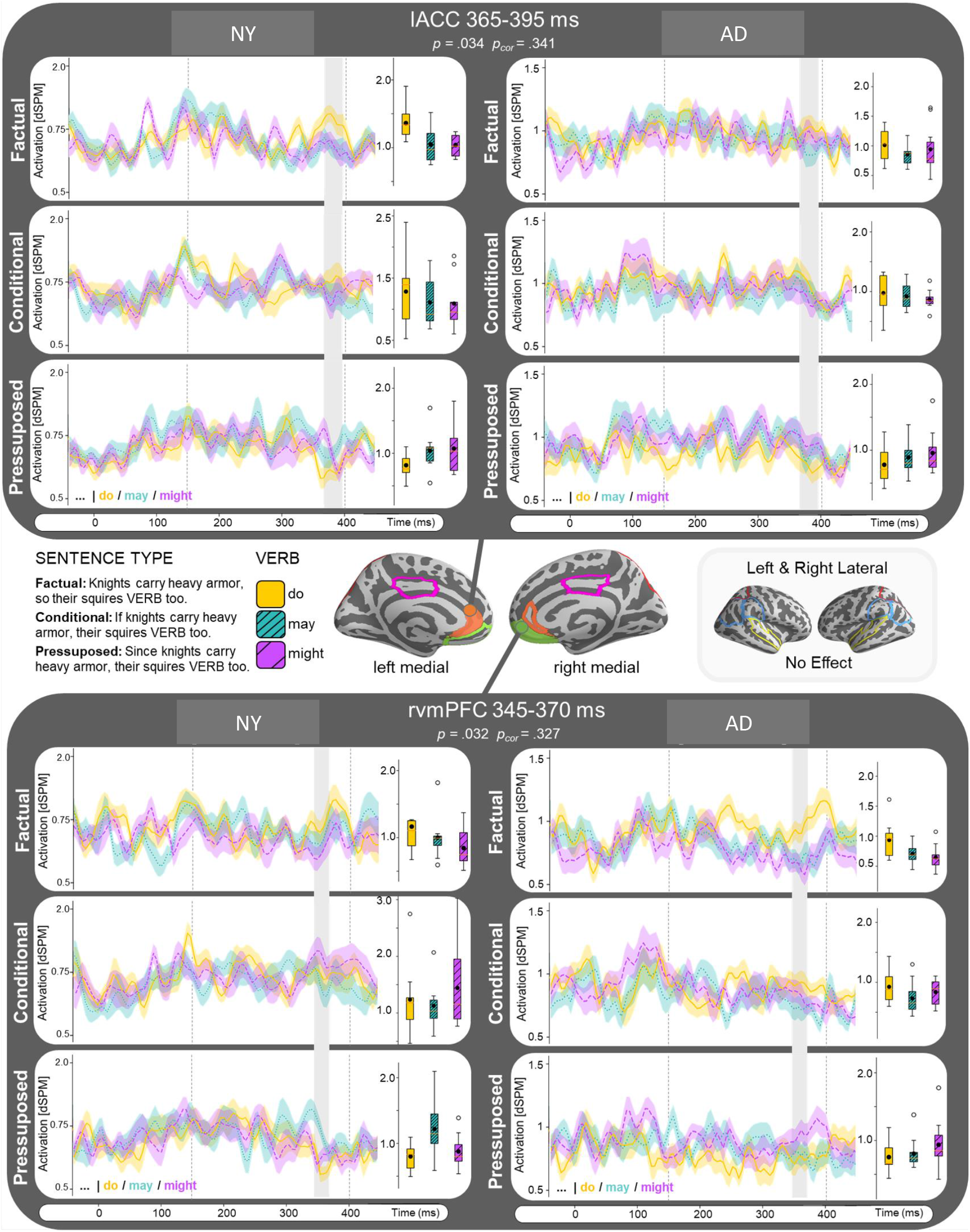

